# ATHENA: A deep learning–based AI for functional prediction of genomic mutations and synergistic vulnerabilities in prostate cancer

**DOI:** 10.1101/2025.11.26.690813

**Authors:** Siyuan Cheng, Xiao Jin, Jiaying Qian, Yuyin Jiang, Isaac Yi Kim, Su Deng, Ping Mu

## Abstract

Identifying functional mutations that drive therapy resistance remains a major challenge in prostate cancer. Large-scale sequencing often produces extensive lists of mutations but provides limited insight into which alterations are functionally relevant. To overcome this gap, we developed **ATHENA** (Attention-based Therapeutic Network Analyzer), a deep learning–based AI framework that predicts the functional impact of genomic mutations and reveals their synergistic vulnerabilities. Integrated with our RNA/DNA-informed variant discovery pipeline **OncoVar**, ATHENA models nonlinear dependencies among mutations to distinguish driver events from passenger variants. Trained on large multi-cohort datasets and interpreted using SHAP analysis, ATHENA not only stratifies patients by clinical outcomes but also predicts which specific mutations alter tumor behavior and therapy response, enabling direct validation through base editing experiments. Applied to prostate cancer progression models, the OncoVar–ATHENA framework identified stage-specific driver signatures across castration-resistant, AR-variant–driven, and metastatic disease, and uncovered cooperative interactions such as SYVN1–STC2 that promote tumor proliferation. By moving beyond simple mutation identification, ATHENA enables functional prediction of genomic interactions. This approach accelerates the discovery of actionable targets and provides a foundation for rational design of next-generation combination therapies in advanced prostate cancer.

## 1. Introduction

Prostate cancer (PCa) is the most frequently diagnosed non-cutaneous malignancy in men and is projected to affect approximately 313,780 men in the United States in 2025^1^. Androgen-deprivation therapy (ADT), together with next-generation androgen receptor (AR) pathway inhibitors such as abiraterone, enzalutamide, and apalutamide, has extended survival in advanced disease. However, resistance ultimately develops in nearly all cases, underscoring the urgent need to identify new biomarkers and therapeutic targets to guide treatment before and after resistance emerges^2,3^.

Molecular profiling has revealed that castration-resistant prostate cancer (CRPC) arises through extensive epigenetic reprogramming, including alterations in DNA (hydroxy)methylation and chromatin-modifier activity^4–14^, as well as through diverse genomic aberrations. Loss-of-function mutations in RB1 and TP53, which occur in 25–60% of metastatic CRPC, promote lineage plasticity and support AR-independent growth^10^. PTEN deletions hyperactivate PI3K–AKT signaling in up to 60% of cases^15,16^, while recurrent defects in DNA damage repair genes such as BRCA1/2, ATM, and CDK12 affect responses to PARP inhibition^17,18^. The AR locus itself is frequently altered, with amplification present in approximately 60% of metastatic tumors and mutations in 15–20%^3,19^. In addition, alterations in transcriptional regulators including HOXB13^20,21^, FOXA1^22,23^, and the TMPRSS2–ERG fusion^24,25^ further reshape the oncogenic landscape.

Despite these advances, stratifying patients based on genomic information remains a major challenge. Most tumors harbor a long tail of rare, often subclonal or heterozygous mutations that are poorly captured by bulk sequencing and remain functionally uncharacterized^26–28^. Moreover, current genomic and prognostic models generally treat mutations as independent variables, overlooking higher-order (epistatic) interactions that may collectively drive therapy resistance. Consequently, reliable tools to identify patients likely to fail AR-targeted therapies or to prioritize rational combination treatments remain limited.

To overcome these challenges, we developed a new artificial intelligence (AI) framework that fundamentally transforms how genomic mutations are interpreted in prostate cancer. This integrated system combines ***OncoVar***, a joint RNA–DNA variant-calling pipeline we newly developed, with ***ATHENA*** (Attention-based Therapeutic Network Analyzer), a deep learning–based AI we designed to predict the functional consequences of individual mutations and reveal their synergistic interactions. OncoVar leverages transcript-level evidence and structure-informed pathogenicity scoring to capture transcriptionally active, low-frequency driver events often missed by conventional methods. ATHENA then models non-linear dependencies among these alterations to distinguish functional from passenger mutations and map combinatorial genomic networks driving disease progression and therapy resistance. Trained on large, multi-cohort datasets and interpreted through explainable AI analysis, ATHENA has already identified three distinct, stage-specific driver signatures corresponding to CRPC, AR-variant–driven CRPC (CRPC-ARv7), and metastatic CRPC (mCRPC), each enriched for biologically and clinically meaningful functional mutations. By directly linking computational prediction to experimental validation through base editing and organoid modeling, ATHENA represents a revolutionary advance that bridges AI-driven genomics with functional cancer biology, accelerating discovery of actionable vulnerabilities and reshaping the precision oncology paradigm in prostate cancer.

## 2. Results

### 2.1. Develop OncoVar for High-confidence Identification of Expression-supported Pathogenic Mutations

To address the long-standing challenge of detecting rare but functionally important genomic alterations often missed by bulk sequencing, we developed OncoVar. It is an integrated computational framework that combines DNA and RNA variant evidence into a single, high-resolution discovery pipeline. Unlike traditional DNA-only approaches, OncoVar is designed to leverage the complementary strengths of paired DNA and RNA sequencing data, providing a more sensitive and biologically grounded view of mutational events. Built within a reproducible nf-core^29^ environment, the system consolidates robust variant-calling algorithms: Mutect2 and HaplotypeCaller through the Sarek workflow for comprehensive somatic and germline discovery from DNA, and HaplotypeCaller through the RNAvar workflow for transcriptionally supported variant detection from RNA^30^. By jointly integrating these parallel outputs, OncoVar enables the identification of low-allele-fraction and transcriptionally active mutations that would otherwise escape detection. This dual-evidence strategy uniquely enriches for variants that are not only present in the tumor genome but also actively expressed, providing a strong support for their biological relevance.

This automated pipeline (Fig. 1A) links data acquisition with parallelized processing and incorporates a central innovation: the integration of AlphaMissense for structure-informed pathogenicity scoring. AlphaMissense, an advanced AI model adapted from DeepMind’s AlphaFold platform, effectively addresses the long-standing difficulty of interpreting variants of unknown significance by combining predicted protein structural context with deep evolutionary conservation features^31^. Through this integration, AlphaMissense provides highly accurate pathogenicity scores for missense mutations, enabling OncoVar to evaluate not only whether a variant is present and expressed, but whether it is structurally and evolutionarily likely to alter protein function. OncoVar systematically incorporates these scores into its final prioritization stage, merging them with joint DNA–RNA evidence in a comprehensive overlap analysis that isolates a uniquely refined set of high-confidence, expression-supported, and structure-informed pathogenic variants. This multilayered approach allows OncoVar to move beyond traditional genomic pipelines by identifying variants with both genomic validity and functional plausibility. As a result, OncoVar delivers a curated catalog of candidate driver mutations that serves as a powerful foundation for downstream modeling with ATHENA, enabling subsequent prediction of mutation function, synergistic interactions, and their contributions to prostate cancer progression and therapeutic resistance.

**Figure 1.**
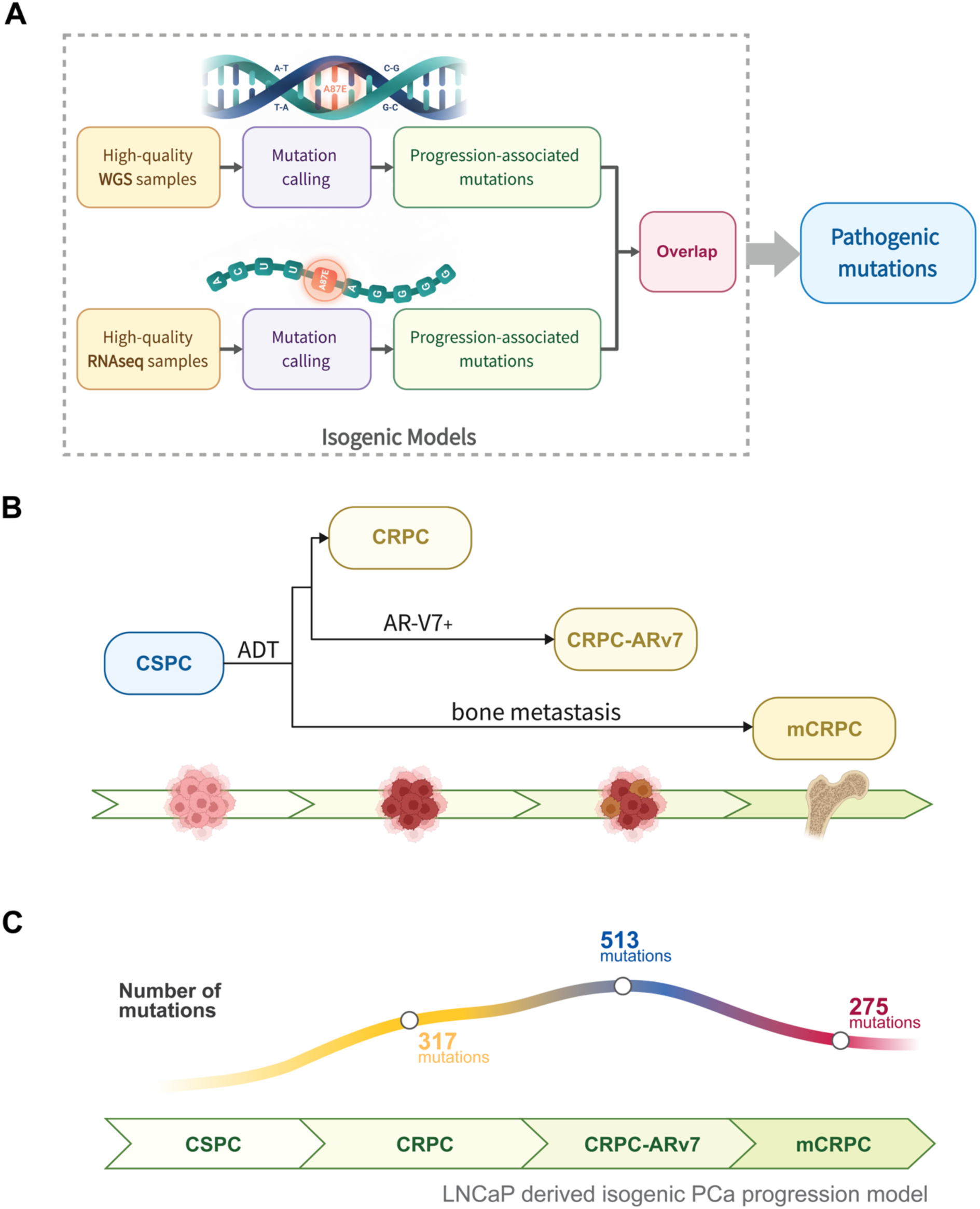
The OncoVar workflow and its application to an isogenic prostate cancer progression model. (A) Overview of the OncoVar workflow, which integrates paired DNA–RNA variant calling, whole-genome validation, and structure-informed pathogenicity scoring to identify high-confidence, expression-supported mutations. (B) Isogenic prostate cancer progression model used to isolate drivers of therapeutic resistance. All cell lines are derived from the castration-sensitive prostate cancer (CSPC) LNCaP parental line and represent distinct clinical stages of disease progression: LNCaP-abl models AR-hypersensitive CRPC, LNCaP95 models ARv7-driven CRPC, and C4-2B represents the aggressive bone-metastatic mCRPC state. (C) Number of CRPC-private, expressed mutations identified by OncoVar across the progression series after computational subtraction of the parental LNCaP genomic background. The results reveal a stepwise accumulation of acquired driver alterations from CSPC to advanced CRPC, followed by a pronounced reduction in the mCRPC stage, consistent with a metastatic bottleneck that selects a refined subset of highly potent driver mutations.

### 2.2. OncoVar identifies treatment-resistant driver mutations in CRPC model

A central challenge in deciphering CRPC is distinguishing clinically relevant driver mutations from the extensive background of passenger alterations. To evaluate our method in a genomically well-defined setting, we applied OncoVar to a high-fidelity isogenic progression model composed of independently generated cell lines derived from the castration-sensitive prostate cancer (CSPC) LNCaP parental line (Fig. 1B). These models recapitulate the clinical spectrum of therapeutic failure. The LNCaP-abl line, established through prolonged androgen-deprivation treatment (ADT), represents the transition from castration-sensitive to castration-resistant disease under ADT selection pressure. The LNCaP95 line, characterized by high expression of the constitutively active ARv7 splice variant, models a more aggressive, AR splice–variant–driven disease state. The C4-2B line, derived through in vivo selection from LNCaP followed by bone metastasis isolation, represents the most advanced metastatic CRPC (mCRPC) phenotype. This shared genomic ancestry provides a precise biological filter that enables the identification of de novo alterations driving progression from CSPC to distinct, clinically lethal CRPC states.

This isogenic system offers an ideal biological framework for applying the OncoVar workflow. Our central hypothesis is that the transition from CSPC to lethal CRPC is driven by the acquisition of rare, transcriptionally active mutations—the exact class of alterations that OncoVar is optimized to detect with high sensitivity. We therefore applied OncoVar across the entire progression series. By computationally subtracting the ancestral genomic profile of the LNCaP parental line, we distilled a high-confidence set of CRPC-specific alterations. These mutations, confirmed to be both genomically present (DNA) and functionally expressed (RNA), represent acquired driver events that contribute to therapeutic resistance and disease progression.

To isolate these events, we applied the integrated RNA–DNA variant-calling framework using approximately twenty high-quality RNA-seq replicates per line from the CTPC project^32^. All RNA-detected variants were first cross-referenced against a foundational whole-genome sequencing (WGS) baseline to confirm their identity as bona fide genomic mutations. We then applied the isogenic filter by removing all variants shared with the CSPC parental genome. To further enrich for robust, line-specific alterations, we constructed a stringent consensus: a mutation was retained only if it was detected in at least 80% of RNA-seq samples for a given resistant line (compared with at least 50% in the parental line). This multi-layered filtering strategy effectively distilled the complex mutational landscape into a high-confidence set of expressed, acquired alterations representing the critical drivers of therapeutic resistance. The resulting landscape (Fig. 1C) revealed a stepwise accumulation of these drivers from CSPC through advanced CRPC, followed by a pronounced reduction in the bone-metastatic mCRPC model. This pattern is consistent with a stringent metastatic bottleneck that selects a refined subset of highly potent driver alterations.

The OncoVar pipeline integrates isogenic subtraction, RNA-based consensus calling, and AI-driven pathogenicity scoring to substantially narrow the initial mutational search space, yielding a refined list of high-confidence CRPC driver mutations (Fig. 2A). Functional annotation of these prioritized alterations revealed the expected distribution of somatic variant consequences: missense mutations formed the predominant class, followed by high-impact stop-gained and splice-site alterations (Fig. 2B). Examination of the genomic distribution across all three resistant states showed that these acquired drivers were broadly dispersed throughout the genome, with focal enrichments rather than restriction to canonical mutational hotspots (Fig. 2C).

**Figure 2.**
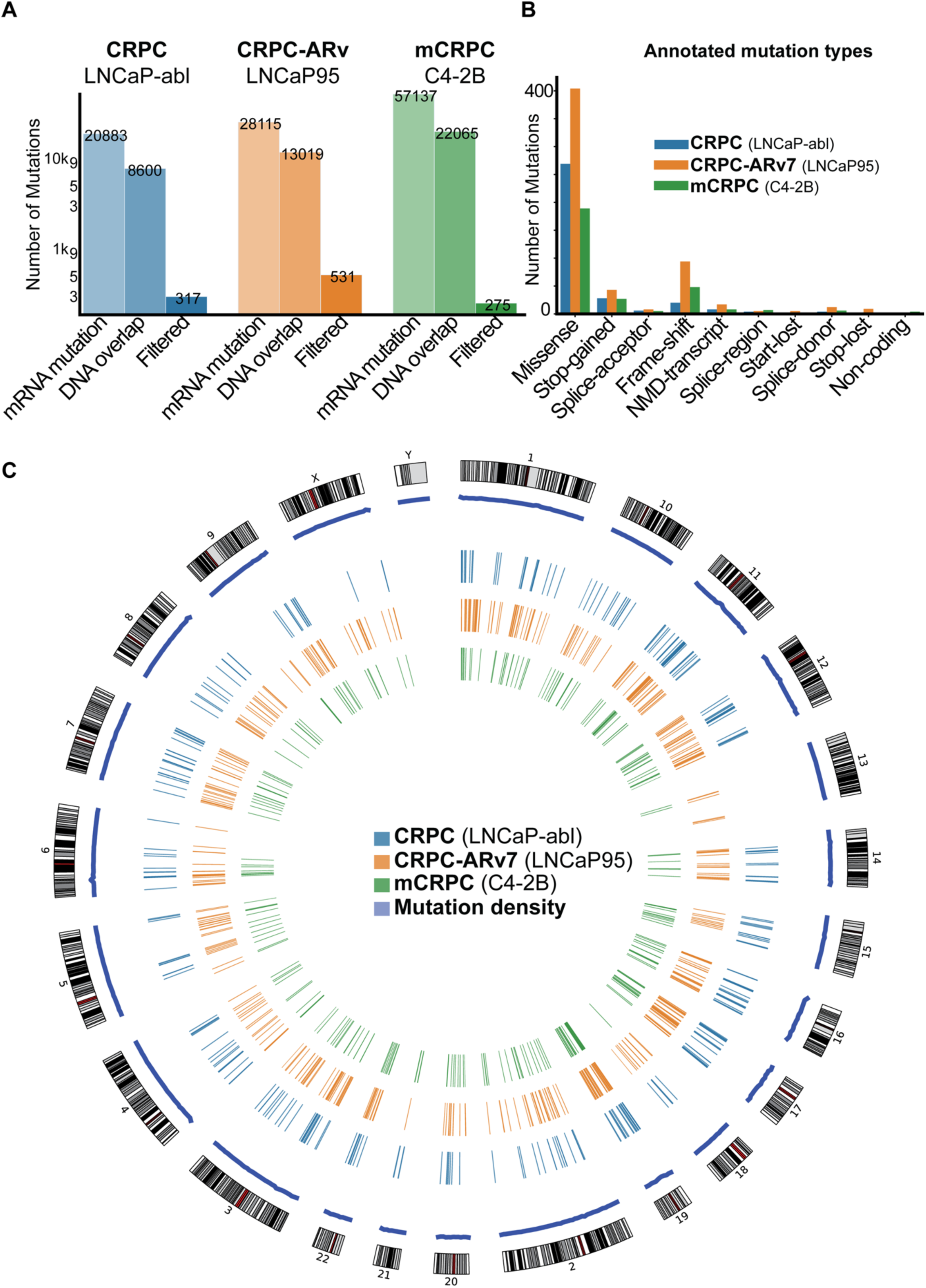
Progressive refinement and genomic distribution of OncoVar-identified CRPC driver mutations. (A) Bar plot showing the number of mutations retained at each filtering step across the three CRPC models, beginning with RNA-seq consensus variant calls and sequentially narrowing through whole-genome sequencing (WGS) cross-referencing and structure-informed pathogenicity scoring, yielding a refined set of high-confidence, expression-supported driver mutations. (B) Distribution of mutation types among the finalized CRPC driver sets for each model, illustrating the predominance of missense alterations followed by high-impact stop-gained and splice-site variants. (C) Circular genome plot showing the chromosomal distribution of the identified driver mutations across all three CRPC stages, revealing that the acquired alterations are widely dispersed throughout the genome with only limited focal enrichments, consistent with resistance evolution being driven by broad genomic diversification rather than a small number of recurrent mutational hotspots.

### 2.3. OncoVar-identified isogenic driver signatures are clinically relevant and prognostic in patient cohorts

Having defined high confidence driver sets in our isogenic CRPC progression model using OncoVar, we next asked whether these signatures capture biologically meaningful patterns in human tumors. To explore this, we analyzed a cohort of 404 metastatic CRPC patients from the SU2C study and mapped each of the three OncoVar driver signatures (CRPC, CRPC ARv7, and mCRPC) onto the corresponding patient tumor genomes. For each patient, the presence of any single-nucleotide variant (SNV) or copy-number variant (CNV) in a gene belonging to a given isogenic signature was counted as a mutation event. This mapping revealed that all three signatures were recurrently altered across the SU2C cohort (Fig. 3A), demonstrating that the stepwise genomic changes captured in our isogenic system are not merely model-specific but are re-enacted in human disease. These results indicate that the isogenic progression model effectively distilled mutational events that mirror real clinical trajectories, supporting the biological fidelity and translational relevance of our OncoVar-derived signatures.

**Figure 3.**
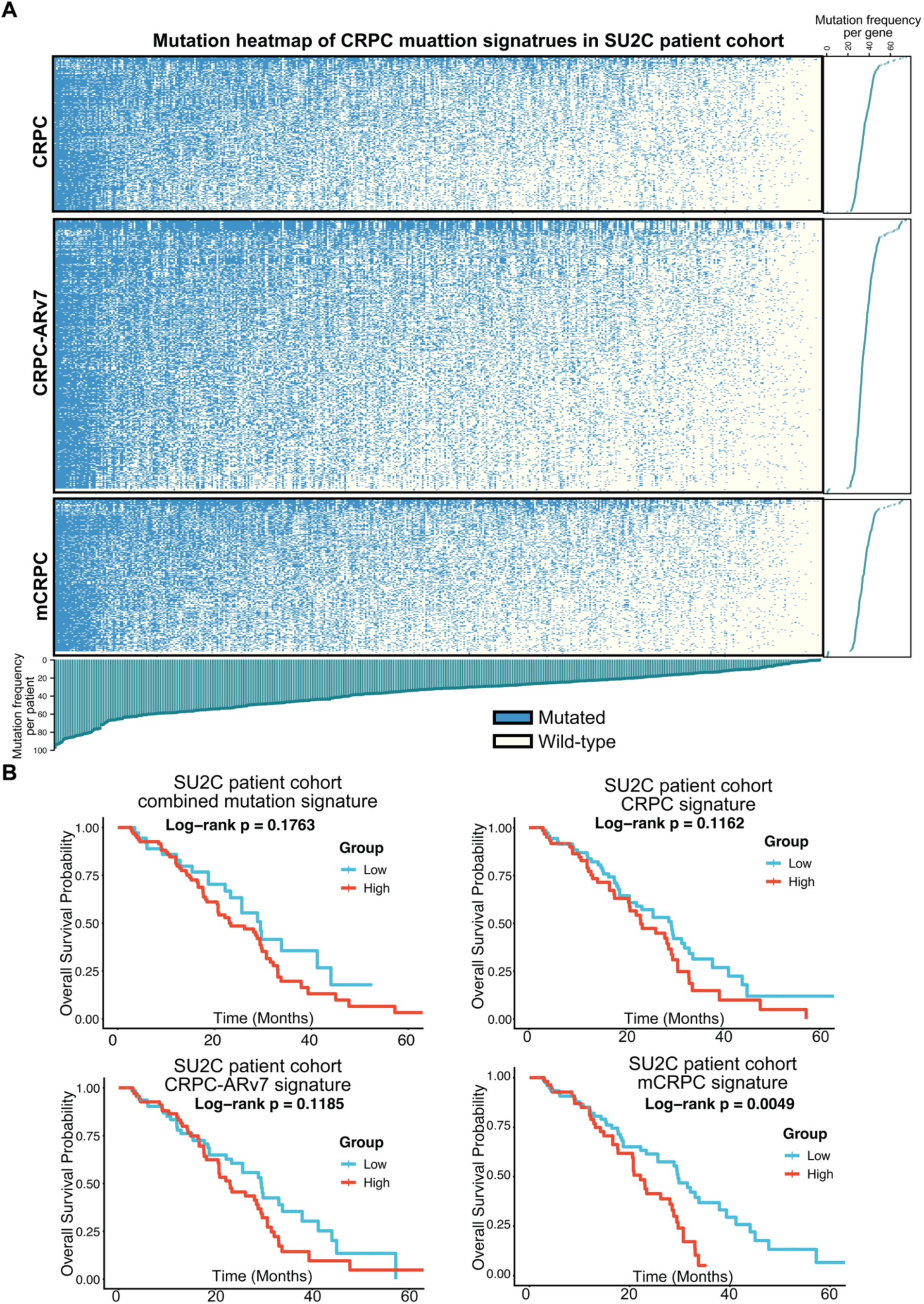
Clinical validation and prognostic significance of isogenic driver signatures in the SU2C patient cohort. (A) Mutation heatmap showing the presence of the three isogenic driver signatures (CRPC, CRPC ARv7, and mCRPC) across 404 metastatic CRPC patients from the SU2C study, where any alteration (SNV or CNV) in a signature gene is annotated as a mutation event. (B) Kaplan–Meier survival analysis evaluating overall survival in patients stratified into high-versus low-mutation-burden groups for each individual signature (CRPC, CRPC-ARv7, and mCRPC) as well as the combined signature.

We next evaluated whether these signatures possess prognostic utility in patients. Kaplan–Meier survival analysis showed that the CRPC and CRPC ARv7 signatures displayed similar risk trends but did not reach statistical significance (Fig. 3B). In contrast, the mCRPC signature exhibited a striking association with clinical outcome: patients whose tumors harbored a high burden of mCRPC-defined driver mutations experienced significantly reduced overall survival (p < 0.05). This finding suggests that the mutations uniquely emerging in the metastatic stage of our isogenic model correspond to the most clinically detrimental alterations in patients, and that the mCRPC signature may define a distinct genomic state linked to aggressive disease biology. Together, these results demonstrate that the OncoVar-derived driver signatures not only recapitulate key features of prostate cancer progression but also carry meaningful prognostic information in patient populations.

### 2.4. Development of the deep learning model ATHENA stratifies metastatic castration-resistant prostate cancer patients by learning non-linear interactions between driver mutations

A major challenge in using mutation signatures as biomarkers or prognostic features is that, although our isogenic system captures a broad landscape of resistance-associated alterations, clinical outcomes are almost certainly driven by combinations of these mutations rather than by individual events considered in isolation. This limitation reflects a broader issue in prostate cancer genomics: most conventional statistical approaches treat mutations as independent variables and are therefore unable to resolve the higher-order, non-linear interactions that likely shape lethal disease progression. These considerations motivated the development of an interaction-aware modeling framework capable of learning how groups of driver mutations jointly contribute to advanced prostate cancer biology.

To overcome the limitations of linear prognostic models and to account for the combinatorial effects of the identified drivers, we developed **ATHENA** (Attention-based THErapeutic Network Analyzer). ATHENA is a multi-branch, attention-based deep learning model designed to identify higher-order combinations of mutations that most strongly influence poor survival, an inference problem beyond the capability of traditional biostatistical approaches. This challenge is further amplified by the fragmented and heterogeneous nature of available genomic datasets, which range from smaller whole-exome cohorts (e.g., SU2C) to large, targeted sequencing panels (e.g., MSK IMPACT). ATHENA directly addresses these issues through a strategy that allows it to distinguish true wild-type states from unprofiled genomic regions, enabling effective integration across datasets with differing genomic coverage.

ATHENA contains three parallel model branches corresponding to the CRPC, CRPC ARv7, and mCRPC driver mutation signatures (Fig. 4A). Each branch incorporates a gated attention module that learns context-dependent attention weights, allowing the model to selectively emphasize the most consequential combinatorial patterns within each signature. The integrated feature representations from these branches are then passed to cohort-specific prediction modules. Training proceeds in two phases: initial pretraining on the large MSK IMPACT cohort to learn broadly applicable patterns, followed by fine-tuning on the SU2C cohort to refine prognostic performance for metastatic CRPC. This combined strategy allows ATHENA to achieve both generalizable robustness and cohort-specific specialization.

**Figure 4.**
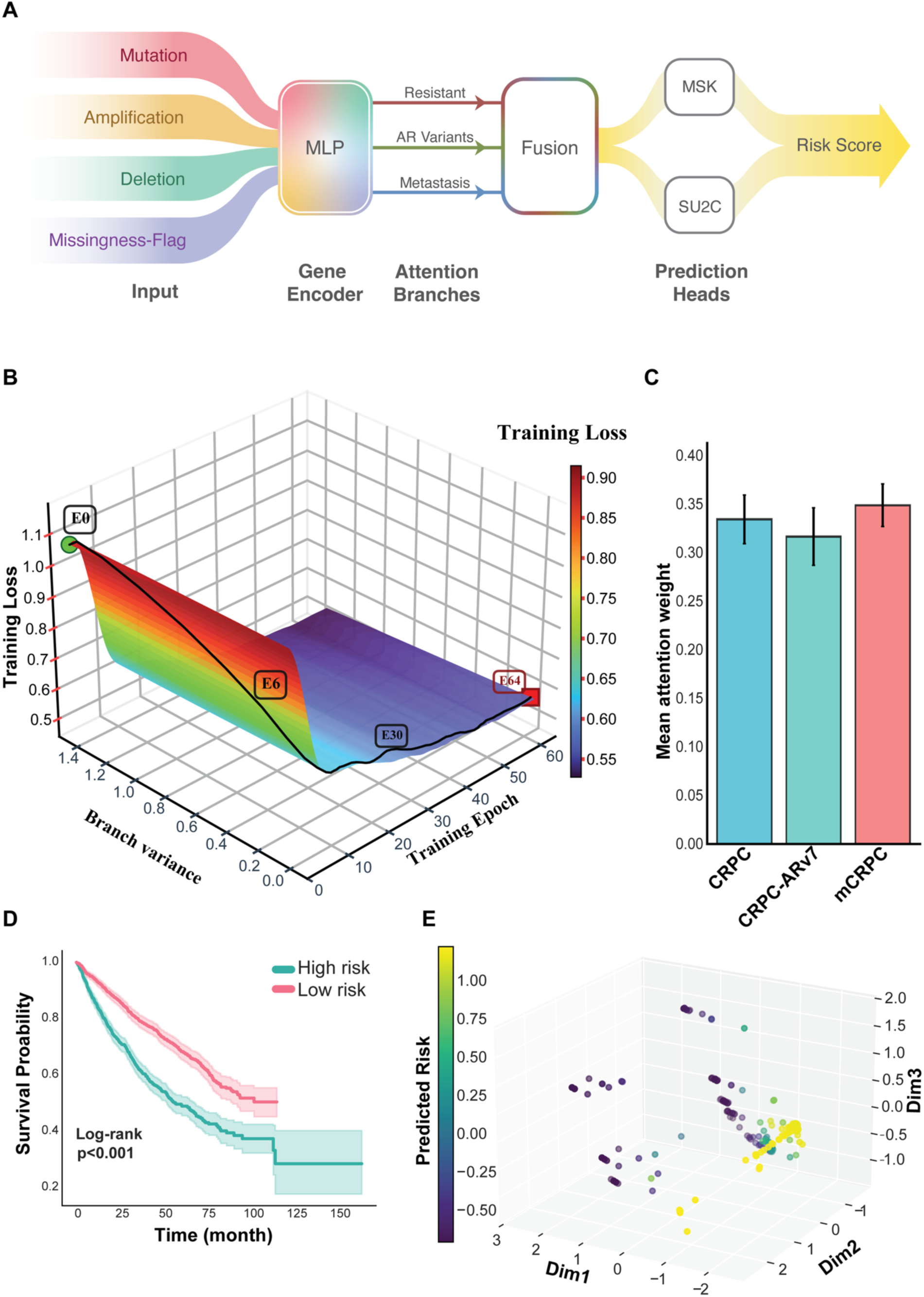
Architecture, training, and performance of the ATHENA deep learning model for prognostic stratification. (A) Schematic overview of the ATHENA (Attention-based Therapeutic Network Analyzer) architecture. The model receives genomic alteration data (mutation, amplification, deletion) together with a learned missingness flag as input. A multilayer perceptron (MLP) gene encoder generates feature embeddings, which are processed by three parallel attention branches corresponding to the CRPC, CRPC-ARv, and mCRPC isogenic driver signatures. The attention-weighted features are then fused and passed to cohort-specific prediction heads (MSK and SU2C) to produce a unified risk score. (B) Three-dimensional visualization of training dynamics showing rapid convergence of training loss across epochs and stabilization of inter-branch variance, indicating a stable and efficient optimization trajectory. (C) Bar plot of the mean attention weights assigned to each of the three isogenic driver signatures by the trained ATHENA model. (D) Kaplan–Meier survival analysis demonstrating ATHENA’s prognostic performance in the combined MSK and SU2C patient cohorts; stratification into high- and low-risk groups based on the model’s risk score reveals a highly significant difference in overall survival (log-rank p < 0.001), with the table below indicating patients at risk at each time point. (E) Three-dimensional UMAP embedding of patient representations learned by ATHENA, where each point corresponds to an individual patient colored by predicted risk, revealing a clear organization along a continuous gradient from low risk (purple) to high risk (yellow).

The model’s training dynamics support its stability and interpretability. The training loss converges quickly, and attention weights across branches stabilize in parallel, indicating that ATHENA consistently learns contributions from each mutation signature (Fig. 4B). Quantifying these weights (Fig. 4C) revealed that all three signatures contribute meaningfully to the final prediction, with the mCRPC signature receiving the highest influence, consistent with its derivation from the most aggressive model.

The critical test of ATHENA’s utility lies in its prognostic performance. Applying the fully trained model to the integrated patient cohort yielded a continuous risk score for each individual. Stratifying patients into high- and low-risk groups based on this score resulted in a striking separation in overall survival (Fig. 4D) (p < 0.001), demonstrating that ATHENA captures clinically relevant interactions missed by mutation-count–based or linear models.

To further understand how ATHENA organizes patient genomes, we visualized the model’s latent space using uniform manifold approximation and projection (UMAP) (Fig. 4E). Patients with low predicted risk (purple) formed distinct clusters from those with high predicted risk (yellow), and a clear gradient from low to high risk emerged across the embedding. This structured separation indicates that ATHENA has learned a biologically coherent representation of metastatic prostate cancer, reflecting the non-linear interplay of driver mutations and their combined effects on prognosis.

### 2.5 ATHENA’s attention mechanism deciphers synergistic driver interactions underlying lethal phenotypes

Having established ATHENA’s robust prognostic performance, we next moved from prediction to mechanistic interpretation. A major limitation of conventional genomic analyses is their inability to resolve higher-order, epistatic interactions among mutations that jointly shape tumor behavior. To uncover these interactions, we used an explainable AI approach based on **SHAP** (SHapley Additive exPlanations). SHAP is a model-agnostic method that assigns each feature, in this case, each gene mutation, a quantitative contribution to the model’s final prediction. Importantly, SHAP can also evaluate pairwise contributions, allowing us to assess how two mutations cooperate to influence the predicted risk score.

We applied SHAP analysis independently to each of ATHENA’s three signature branches and generated mutational synergy heatmaps that reveal the cooperative networks most strongly associated with poor patient survival (Fig. 5). This interaction landscape highlights a central strength of ATHENA: rather than identifying mutations in isolation, it learns how combinations of mutations work together to drive lethal phenotypes.

**Figure 5.**
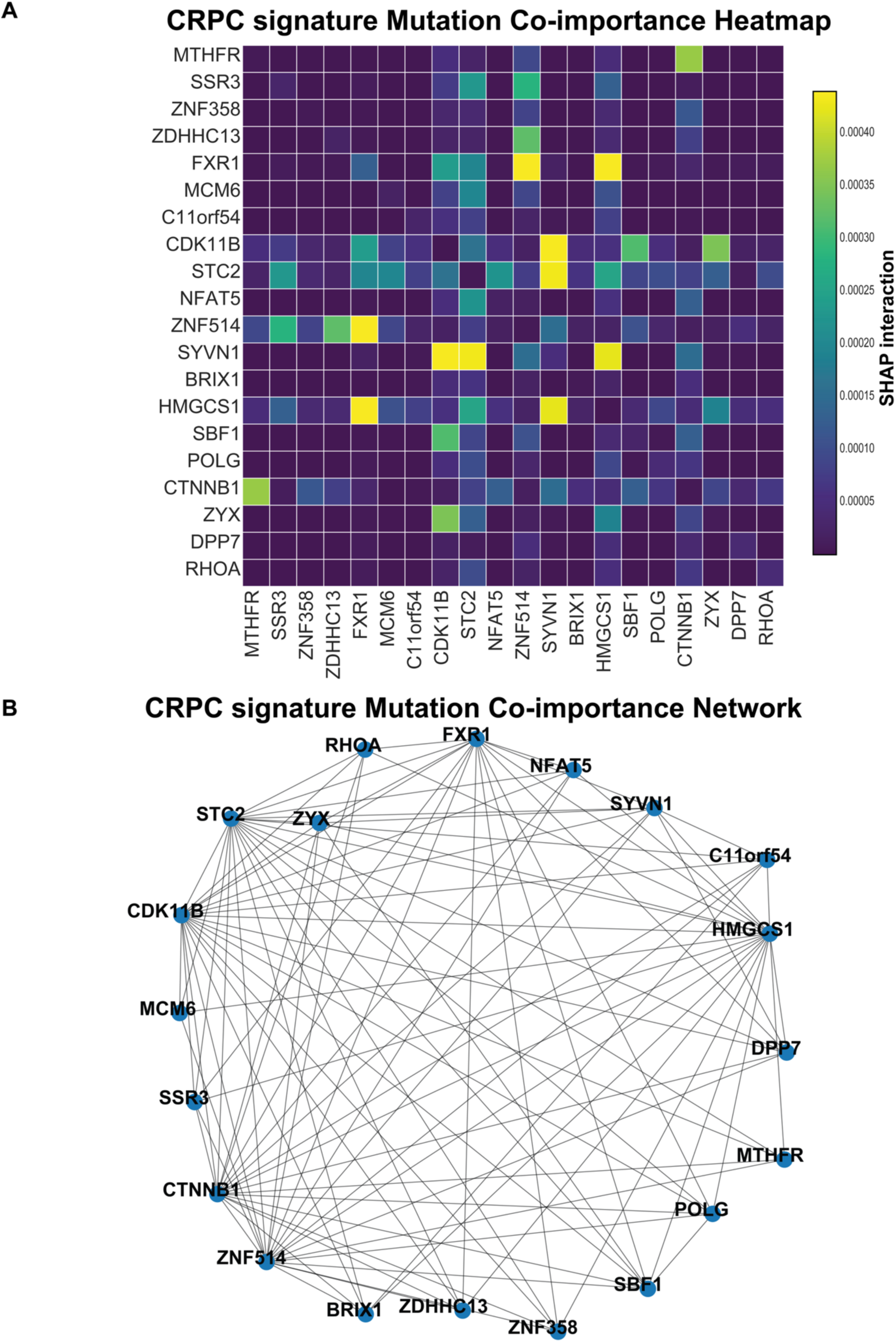
Synergistic interactions among gene mutations in the CRPC driver signature. (A) Heatmap of pairwise SHAP interaction values for all genes within the CRPC signature, with genes shown along both axes. Each cell represents the magnitude of the interaction between two mutations, with brighter colors (yellow) indicating strong synergistic or co-dependent effects, where the combined influence of two genes on the model’s prediction exceeds the sum of their individual contributions. (B) Network representation of the same CRPC co-importance interactions, where each gene is depicted as a node and edges represent interaction strength. This visualization highlights the dense and interconnected cooperative landscape within the CRPC driver signature, revealing clusters of mutations that jointly contribute to aggressive disease behavior.

Having established ATHENA’s robust prognostic performance, we next moved from prediction to mechanistic interpretation. A major limitation of conventional genomic analyses is their inability to resolve higher-order, epistatic interactions among mutations that jointly shape tumor behavior. ATHENA explicitly addresses this challenge. Using SHAP (SHapley Additive exPlanations), we systematically quantified how pairwise combinations of somatic mutations synergize to influence the model’s final risk score. SHAP analysis was performed independently for each of ATHENA’s three signature branches, and the resulting synergy heatmaps revealed cooperative mutational networks most strongly associated with poor patient survival (Fig. 5). This interaction landscape highlights a central strength of ATHENA: rather than simply identifying which mutations are present, it learns how mutations work together to drive lethal phenotypes.

Strikingly, instead of the diffuse or noisy patterns often produced by high-dimensional “black-box” models, the synergy heatmaps revealed well-defined interaction clusters among genes with established or emerging relevance in CRPC biology. Prominent hotspots included genes involved in protein-folding stress responses (such as SYVN1, an ER-resident E3 ubiquitin ligase)^33^, sterol and lipid biosynthesis (such as HMGCS1)^34^, and cell-cycle regulation (such as CDK11B)^35^, all pathways known to support tumor adaptation to androgen receptor (AR) blockade. SYVN1 exhibited some of the strongest synergy signals across the entire matrix. Its coordinated co-important with STC2 and CDK11B suggest that the ER-associated degradation pathway may cooperate with metabolism and mitotic machinery to maintain proteostasis during AR deprivation^34–37^.

Together, these findings demonstrate that ATHENA functions not only as a prognostic model but also as a mechanistic engine capable of uncovering multi-gene cooperative modules that likely represent functional dependencies in therapy-resistant disease. By mapping these epistatic networks, ATHENA provides a biologically interpretable foundation for biomarker discovery and the prioritization of therapeutic targets in advanced prostate cancer.

## Discussion

In this study, we address two of the most formidable challenges in cancer genomics: extracting meaningful signal from the long tail of mutations, and resolving the complex, non-linear interactions among mutations that collectively drive therapeutic resistance. We present an integrated framework that combines a high-fidelity variant discovery pipeline, OncoVar, with a biologically informed deep learning model, ATHENA. Together, these tools distill sparse and heterogeneous genomic data from CRPC into functional, stage-specific driver signatures, CRPC, CRPC-ARv, and mCRPC, enabling the development of a highly accurate prognostic algorithm and, importantly, providing a platform for generating mechanistic hypotheses about the cooperative molecular networks that give rise to lethal disease states. For biologists and prostate cancer researchers, this framework offers a powerful means to systematically move from lists of mutations to functionally grounded hypotheses about how genomic alterations shape resistance trajectories.

The first major advance of this work is the development of OncoVar, an integrated variant-discovery framework designed to overcome fundamental limitations of conventional genomic analysis. Standard approaches are obscured by overwhelming passenger noise, have limited power to detect low-allele-fraction mutations, and implicitly assume that any DNA-level alteration is biologically active. However, many genomic alterations are transcribed at very low levels and are therefore unlikely to have functional relevance^38^. OncoVar resolves these barriers by requiring transcriptional evidence from RNA sequencing, thereby restricting analysis to alterations with the potential for biological impact. Genomic validation using whole-genome sequencing eliminates spurious transcriptional artifacts, and incorporation of AlphaMissense pathogenicity scoring provides a rigorous means of prioritizing variants of unknown significance that would otherwise remain intractable. This design yields a refined set of transcriptionally active, structure-informed driver mutations that serve as a high-fidelity foundation for downstream modeling. Importantly, these filtered alterations represent an enriched pool of candidates for functional studies using base editing, CRISPR knock-in models, and organoid systems, bridging computational predictions with experimental validation in ways directly relevant to prostate cancer biology.

Building on this platform, our second major innovation, ATHENA, introduces a conceptual shift from both generic “black-box’’ deep learning models and oversimplified linear statistical methods. While pan-cancer foundation models leverage immense training datasets, their relevance to a narrowly defined biological question such as CRPC resistance is inherently constrained. Subtle, CRPC-specific combinatorial patterns are diluted by dominant drivers from unrelated malignancies, and such models lack an inductive bias that reflects the sequential and modular nature of CRPC evolution. ATHENA is deliberately engineered as an alternative to this paradigm. Its multi-branch, attention-based architecture mirrors the modular driver signatures uncovered by OncoVar, enabling the model to learn the context-dependent relevance of each resistance phenotype. For prostate cancer scientists, this means ATHENA can identify not only whether a mutation matters, but *why* it matters in the specific setting of AR blockade, lineage plasticity, or metastatic evolution.

ATHENA demonstrates strong prognostic performance (C-index > 0.68) across large and diverse patient cohorts. Yet its most transformative value lies in its interpretability. The emergence of a coherent risk continuum in its latent space, together with the use of SHAP analysis to interrogate its internal logic, positions ATHENA as a mechanistic discovery engine. It reveals specific synergistic mutation pairs, such as SYVN1–STC2 in CRPC, that appear to underlie lethal phenotypes. These context-specific interactions, inaccessible to linear models or mutation-count approaches, highlight biological dependencies that may represent new vulnerabilities.

The translational implications of this framework are substantial. The ATHENA risk score provides a prognostic tool that outperforms stratification based on single-gene biomarkers or raw mutation burden, enabling identification of high-risk patients who may benefit from treatment intensification. Moreover, by pinpointing synergistic driver pairs, our approach offers a rational foundation for designing mechanism-guided combination therapies. Co-targeting pathways implicated by high-impact SHAP pairs may offer a promising strategy to overcome resistance in settings where single-agent therapies have limited efficacy. For clinicians and translational researchers, this provides a mechanistically informed rationale for selecting therapeutic combinations tailored to patient-specific mutational landscapes.

Beyond therapy-resistant prostate cancer, the design philosophy underlying ATHENA offers a generalizable blueprint for integrating high-dimensional biological data with clinically meaningful interpretation. The paradigm of (1) using domain-specific knowledge to isolate and structure biologically distinct modules, (2) constructing model architectures that reflect this modularity, and (3) employing interpretable AI methods to generate new, testable hypotheses has broad applicability across complex diseases. In conclusion, this work provides not only a new paradigm for patient stratification in prostate cancer but also a path forward for developing interpretable, biologically grounded AI systems capable of deciphering the combinatorial logic of disease across all of medicine. For the prostate cancer research community, OncoVar and ATHENA together create a scalable, mechanistically informed platform that accelerates the functional discovery of driver mutations and opens new avenues for understanding, and targeting, the molecular determinants of therapeutic resistance.

## 3. Methods

### 3.1 Patient Cohorts and Genomic Data

Genomic and clinical data were aggregated from two publicly available metastatic prostate cancer cohorts: the Stand Up To Cancer (SU2C) 2019 cohort^39^ (prad_su2c_2019) and the Memorial Sloan Kettering (MSK) 2024 cohort^40^ (prostate_msk_2024). Data, including somatic mutations, copy number alterations (CNA), and clinical outcomes (overall survival time and status), were acquired from the cBioPortal for a total of 2,426 patients^41^. Each patient’s genomic profile was encoded as a tensor of shape (number genes, 4), representing mutation, amplification (alteration >= 2), deletion (alteration = -2), and a critical missingness flag. This flag explicitly marked genes not assayed in eachpatient’s sequencing panel (e.g., MSK-IMPACT panels), enabling the model to distinguish unassayed genes from true wild-type states.

### 3.2 The OncoVar Workflow for Driver Mutation Discovery

The driver discovery was based on a high-fidelity isogenic progression model derived from the parental LNCaP (CSPC) cell line. Derivative lines representing distinct stages of resistance were used: LNCaP-abl (CRPC), LNCaP95 (CRPC-ARv), and C4-2B (mCRPC). High-coverage whole-genome sequencing (WGS) and deep RNA sequencing (RNA-seq, ∼20 replicates per line) data were generated for all lines^32,42^.

To achieve high-confidence variant detection, we implemented a joint WGS and RNA-seq strategy. For WGS, reads were aligned to the GRCh38 reference genome using BWA-MEM (v0.7.17), processed according to GATK4 (v4.2.6.1) best practices, and variants were called using both GATK Mutect2 and FreeBayes (v1.3.6). For RNA-seq, we utilized the nf-core/rnavar pipeline, which implements GATK4 best practices for RNA data, including STAR alignment (v2.7.9a) and HaplotypeCaller for variant detection.

To isolate acquired drivers of resistance, we performed a multi-step filtering process. First, we computationally subtracted all variants present in the parental LNCaP WGS data from the derivative line call sets. Next, to ensure variants were robustly expressed, a consensus was required: a variant was retained only if detected in ≥80% of RNA-seq samples for a given resistant line. Finally, all high-confidence variants, confirmed in both WGS and RNA-seq, underwent functional annotation using Ensembl VEP (v104.3). Missense variants were further prioritized based on their predicted pathogenicity using AlphaMissense, yielding three distinct, stage-specific driver signatures.

### 3.3. Survival analysis and data retrieval

Patient data preprocessing was performed to construct binary mutation matrices from the cBioPortal datasets described in Section 4.1. Mutation data were filtered to retain only non-synonymous variants (missense, nonsense, frameshift, in-frame insertions/deletions, and splice site mutations). For each cohort, a binary mutation matrix was created with dimensions (number of patients × number of genes), where entries of 1 indicated the presence of a mutation in that gene for that patient, and 0 indicated wild-type status. Copy number alterations were encoded separately, with amplifications represented as +2 and deletions as -2. Patient identifiers were standardized across mutation and clinical datasets to enable merging.

Three primary analyses were performed on the processed data. Mutation burden histograms were generated using ggplot2 (v3.4.0) with 30 bins to visualize the distribution of total mutated genes per patient across both cohorts (SU2C and MSK). Heatmap visualizations were constructed using the ComplexHeatmap package (v2.24.1) to display mutation patterns across patients and genes. The binary mutation matrix was subset to include only genes from the three isogenic driver signatures (CRPC, CRPC-ARv7, and mCRPC). Patients and genes were hierarchically clustered using Euclidean distance and complete linkage, with the heatmap annotated with patient mutation burden (row annotations) and gene mutation frequency (column annotations). Survival analysis was performed to assess the prognostic significance of mutation burden, with patients stratified into high and low mutation burden groups based on the median mutation load. Kaplan-Meier survival curves were generated and visualized using the survival (v3.8-3) and survminer (v0.4.9) packages, with the log-rank test used to assess statistical significance of differences in overall survival between groups.

### 3.4 The ATHENA Deep Learning Model

A two-phase transfer learning strategy was employed. In Phase 1, the main prediction head was trained on the MSK cohort. In Phase 2, the specialized SU2C head was fine-tuned on the SU2C cohort, with the main model weights frozen. The training objective was a composite loss function combining the Cox partial likelihood loss with several regularization terms: a concordance loss (C-index), a branch diversity loss to prevent feature collapse, an attention entropy loss to encourage balanced branch usage, and a contrastive separation loss to maximize risk differences between event and censored patients. The AdamW optimizer was used with learning rates of 3e-4 (Phase 1) and 2e-4 (Phase 2). Early stopping based on validation of C-index was used to prevent overfitting.

Model performance was primarily evaluated using the concordance index (C-index) on held-out validation sets. Kaplan-Meier survival analysis was used to visualize the prognostic separation of risk groups defined by the model’s output. All data splits were stratified by event status for balanced evaluation.

To move from prediction to mechanistic insight, we employed SHAP (SHapley Additive exPlanations). We calculated pairwise SHAP interaction values for mutations within each of the three signature branches. This allowed us to quantify the synergistic effect of mutational pairs on the final risk score, thereby revealing the cooperative networks learned by the model.

Analyses were performed in Python (v3.8). The deep learning model was built using PyTorch (v1.10.0). Survival analyses utilized the lifelines (v0.27.0) package, and SHAP analysis used the shap (v0.44.0) package. Data processing and visualization relied on numpy, pandas, scikit-learn, matplotlib, and seaborn. The code and data used in this study will be made available upon publication.

## 3.5 Statistical analysis

Statistical analyses were performed in R (v4.3.0). Data manning utilized the tidyverse collection of packages (v2.0.0), including dplyr for data filtering and transformation, and tidyr for matrix reshaping. For survival analysis, the log-rank test was used to compare survival distributions between groups, with statistical significance assessed at α = 0.05. Hierarchical clustering for heatmap visualization employed Euclidean distance metrics with complete linkage. All statistical analyses and visualizations were implemented using base R functions and the aforementioned specialized packages. All code and analysis scripts are available upon request.

## Acknowledgments

This work was supported or partially supported by: National Cancer Institute/National Institutes of Health: R37CA258730, R01CA288820, R01CA292949 P. Mu; Prostate Cancer Foundation: 25CHAL05, 17YOUN12 P. Mu, Yale Cancer Center CCSG Pilot Grant. P30CA016359 P. Mu. and I.Y.K.

## CRediT author statement

**Siyuan Cheng**: Conceptualization, Methodology, Software, Data curation, Writing-Original draft preparation; **Xiao Jin**: Formal Analysis, Data curation, Visualization; **Jiaying Qian:** Investigation, Data Curation, Visualization; **Yuyin Jiang**: Writing-Reviewing and Editing; **Isaac Yi Kim**: Supervision; **Su Deng**: Validation, Supervision; ***Ping Mu***: Supervision, Writing-Reviewing and Editing.

## Conflict of Interest Statement

P.M. served as a scientific consultant to Accutar Biotechnology, Inc. No other authors have COI to disclose.

